# Modelling the Effect of Allopregnanolone on the Resolution of Spike-Wave Discharges

**DOI:** 10.1101/2023.07.06.547738

**Authors:** Maliha Ahmed, Sue Ann Campbell

## Abstract

**Objective:** Childhood absence epilepsy (CAE) is a paediatric generalized epilepsy disorder with a confounding feature of resolving in adolescence in a majority of cases. In this study, we modelled how the small-scale (synapse-level) effect of progesterone metabolite allopregnanolone induces a large-scale (network-level) effect on a thalamocortical circuit associated with this disorder. In particular, our goal was to understand the role of sex steroid hormones in the spontaneous remission of CAE.

**Methods:** The conductance-based computational model consisted of single-compartment cortical pyramidal, cortical interneurons, thalamic reticular and thalamocortical relay neurons, each described by a set of ordinary differential equations. Excitatory and inhibitory synapses were mediated by AMPA, GABAa and GABAb receptors. The model was implemented using the NetPyne modelling tool and the NEURON simulator.

**Results:** The action of allopregnanolone on individual GABAa-receptor mediated synapses has an ameliorating effect on spike-wave discharges (SWDs) associated with absence seizures. This effect is region-specific and most significant in the thalamus, particularly the synapses between thalamic reticular neurons.

**Significance:** The remedying effect of allopregnanolone on SWDs may possibly be true only for individuals that are predisposed to remission due to intrinsic connectivity differences or differences in tonic inhibition. These results are a useful first-step and prescribe directions for further investigation into the role of ALLO together with these differences to distinguish between models for CAE-remitting and non-remitting individuals.

## 1 Introduction

Childhood absence epilepsy (CAE) is an idiopathic generalized epilepsy disorder which affects children between the ages of 4-12 years (Islam & Abdullah, 2014; Wirrell, Camfield, Camfield, Gordon, & Dooley, 1996). It is characterized by sudden brief periods of impaired consciousness occurring several times a day. Unlike some other epilepsy types, CAE is more common in girls than boys (Crunelli & Leresche, 2002). One of the most confounding features of this disease is its age-dependent expression, and the ability to spontaneously resolve in adolescence, with remission occurring in about 70-80% of cases, while in others it can progress into more severe types of epilepsy (Wirrell et al., 1996). There are several hypothesized mechanisms involved in remission, including antiepileptic drugs, differences in circuitry arrangements and genetic predisposition or dynamic epigenetic processes. However, there remains an inadequate understanding of some of these factors that can inform early intervention practices.

Clinically, CAE is diagnosed using an electroencephalogram (EEG) where recordings exhibit bilateral synchronous 2.5-4 Hz spike-wave discharges (SWDs) (Wirrell et al., 1996). The recorded EEG can thus be used as an indicator for underlying brain activity and neuronal networks that may be involved in giving rise to such oscillations. Based on many years of amounting EEG data, the thalamocortical circuit consisting of pyramidal neurons in the cortex, thalamic relay neurons and thalamic reticular neurons in the thalamus are considered to be important in the generation of normal 7-14 Hz sleep spindles as well as in the pathophysiology of absence seizures (Kim et al., 2020). According to the cortical focus theory, the cortex is considered as the driving source for the origin of SWDs, with a cortical focus initiating rhythmic discharges followed by propagation towards generalized activity through intracortical pathways and the thalamus, thus forming an integrated network (Meeren, van Luijtelaar, da Silva, & Coenen, 2005).

There is a considerable amount of evidence towards a strong contribution to the development of SWDs by genes encoding receptors for the inhibitory neurotransmitter *γ*-Aminobutyric acid (GABA), as well as ion channels including calcium channels, sodium channels, and potassium channels (Kole, Bräuer, & Stuart, 2007; Ludwig et al., 2003; Ogiwara et al., 2018; Oliva et al., 2014; Papale et al., 2009). Here we focus on the effect of genes encoding T-type calcium channels and GABAa receptors, which we describe in more detail below.

T-type calcium channels (most prominently Cav3.1, Cav3.2 and Cav3.3) are known to play a key role in the generation and maintenance of spike-wave discharges as they are fundamental in thalamic burst firing (Chen, Parker, & Wang, 2014). Cav3.2 channels encoded by the CACNA1H gene are mainly expressed in thalamic reticular neurons and in some pyramidal neurons in the cortex (Chen et al., 2014). There is a spectrum of CACNA1H variants linked to CAE which results in enhanced T-type calcium current and consequently alters the firing of thalamic neurons to generate SWDs (Chourasia, Ossó-Rivera, Ghosh, Von Allmen, & Koenig, 2019). For instance, the P640L variant of Cav3.2 induces changes in the biophysical properties of the channel including altered current (in)activation time constants, current amplitudes, as well as shifts in voltage dependence on (in)activation (Glauser et al., 2017).

Pharmacological data has shown that inhibitory and excitatory neurotransmissions are crucial in the development and control of absence seizures (Contreras & Steriade, 1995; Cope et al., 2009; Tan et al., 2007). One of the most prevalent factors in the alteration of SWDs is the function of GABA. The intrinsic properties of GABAa receptors (GABAaRs) may vary depending on different subunit compositions. Receptors containing *α*1-3, *β*1-3 and *γ*2 subunits are more commonly localized in postsynaptic sites, and receptors predominantly composed of *α*1*β*2*γ*2 subunits comprise approximately 43-60% of all GABAaRs in the adult brain (Wallace et al., 2001; Yalçın, 2012). Mutations in the GABRG2 gene which encodes the *γ*2 subunit of the GABAaR is most strongly associated with CAE, while variants of GABRA1, GABRB2 and GABRB3 genes are weakly implicated as reported in some studies (Fu, Wang, Kang, & Mu, 2022; Yalçın, 2012). For instance, a GABRG2 heterozygous missense variant R82Q as well as the R43Q mutation were reported to reduce *γ*2 surface expression due to abnormal trafficking resulting in reduced GABAaR currents, reduced surface expression of GABAaRs in pyramidal neurons in the cortex, as well as reduction in miniature inhibitory postsynaptic currents in layer 2/3 cortical neurons (Currie et al., 2017; Macdonald, Kang, & Gallagher, 2010; Marini et al., 2003). Thus, the overall effect of GABAaR subunit gene mutations is disinhibition causing abnormality in cortical network function.

Furthermore, given that remission often occurs at adolescence (a time of onset of puberty), the association of sex hormones with CAE is also suspected (van Luijtelaar et al., 2001). This association is further supported by the phenomenon of catamenial exacerbation of epileptic seizures. Catamenial epilepsy refers to the periodicity of seizure exacerbation that aligns with the cyclical changes in circulating levels of ovarian steroid hormones during the menstrual cycle (Foldvary-Schaefer & Falcone, 2003). The cyclical variation of steroid serum levels and its positive correlation with seizure frequency implies a neuroactive role of sex steroid hormones.

Progesterone is one of the major steroid hormones that impacts sexual development during puberty and is associated with reproduction. It can be synthesized locally in the central and peripheral nervous system or derived from the gonads or adrenal gland (Frye, Koonce, & Walf, 2014; Schumacher et al., 2012). The metabolism of progesterone involves the production of allopregnanolone (ALLO), a neuroactive steroid. Neurosteroids such as allopregnanolone, may be synthesized de novo in the nervous system or accumulate in the brain through the bloodstream after metabolism of adrenal or gonadal steroids (Frye et al., 2014). They rapidly modulate neuronal excitability through both genomic (i.e., regulation of gene expression) and non-genomic mechanisms (i.e., modulation of neurotransmitter-gated ion channels, and activation of signalling cascades) (Brinton et al., 2008). There is substantial evidence from both clinical studies as well as animal experiments on the effect of progesterone (and allopregnanolone) on human and animal EEG (Wheless & Kim, 2002). Administration of progesterone has been shown to result in an exacerbation of absence seizures. (Budziszewska, Van Luijtelaar, Coenen, Leśkiewicz, & Lasoń, 1999; van Luijtelaar et al., 2001). Allopregnanolone is a positive allosteric modulator of the GABAaR. In particular, it increases the probability that an agonist will bind to the receptor as well as the receptor efficacy (Bromfield, Cavazos, & Sirven, 2006; Majewska, Harrison, Schwartz, Barker, & Paul, 1986). The overall effect is to enhance and prolong the action of GABA, resulting in enhanced inhibitory connections.

It is certainly understood that childhood absence epilepsy is a multifactorial disorder which poses many challenges, but also unique opportunities for exploration, when it comes to modelling. The objective of this study was to use a well-established thalamocortical model associated with CAE to explore the role of allopregnanolone in the eventual resolution of absence seizures, as observed in most cases. In particular, our goal was to gain insight on factors that may predispose some individuals to experience remission or as we have termed it, ‘resolving CAE’. This study also explored how the effect of allopregnanolone differs depending on the genetic factors associated with CAE and the consequent ability of the network to transition between a healthy state and diseased state through variation of model parameters.

## 2 Methods

Our thalamocortical model is based on the previous work by Knox et al. (Knox, Glauser, Tenney, Lytton, & Holland, 2018) in which a well-established thalamocortical model by Alain Destexhe (Destexhe, 1998) was used to study the contribution of the T-type calcium channel in thalamic reticular neurons combined with cortical excitability in the pathogenesis and treatment response in CAE.

The model, a 400-cell thalamocortical network, consists of single-compartment neuron models of four types: cortical pyramidal cells (PY), cortical interneurons (IN), thalamic reticular cells (RE) and thalamocortical cells (TC). The membrane potential of each cell type is described by the following equations:

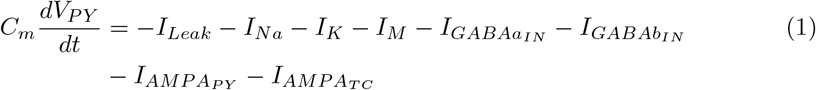

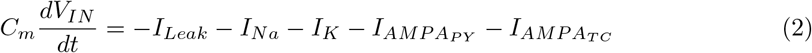

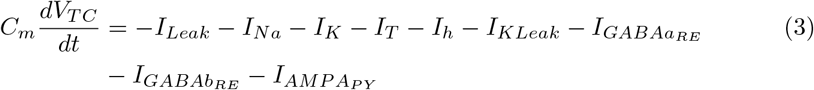

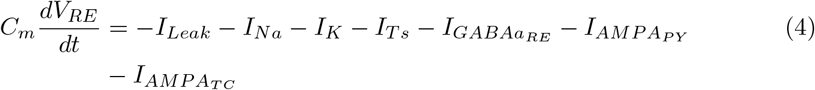

where *V*_*i*_ is the membrane potential for *i* = PY, IN, TC, RE, *C*_*m*_ = 1*µF*/*cm*^2^ is the specific capacity of the membrane. All intrinsic currents follow the same formalism as in Equation (5), 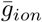 where represents the maximal conductance, and *m* and *h* represent the voltage gating of the ion channels. All synaptic currents follow Equation (6) with AMPA/GABAa- and GABAb-mediated synapses being described appropriately by the synaptic conductance function, *s*(*t*), and 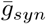 denotes the maximal conductance. All of the cell types have *Na*^+^ and *K*^+^ currents as described in Traub et al. (Traub, Wong, Miles, & Michelson, 1991). The IN cells do not have any additional currents, while the PY cells have one additional slow *K*^+^ current, *I*_*M*_, which contributes to spike frequency adaptation behaviour. Both the thalamic cells exhibit bursting behaviour due to the presence of a low-threshold *Ca*^2+^ current, *I*_*T*_, with slower kinetics in the RE cells. In addition to this, TC cells also have a hyperpolarization-activated current, *I*_*h*_. The kinetics of all the currents are as described in Destexhe et al. (Destexhe, Contreras, & Steriade, 1998), leading to models of the form:

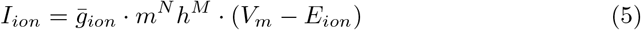

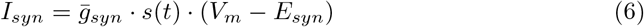

Cortical cells represent layer 6 of the cerebral cortex and have excitatory synapses formed with ascending thalamic axons, creating an excitatory feedback loop. The network is made up of 100 cells of each type and each layer of cells is arranged in one dimension as shown in Figure 1 with connectivity and synapses varying between cell types as described. All excitatory connections in the network are mediated by AMPA receptors, and inhibitory connections are mediated by either just GABAa or a combination of GABAa and GABAb receptors. Within the thalamus and the cortex, each neuron has connections with 11 postsynaptic neurons, while between the thalamus and cortex, each presynaptic neuron has connections with 21 postsynaptic neurons, as illustrated in Figure 1. The synaptic conductance values were chosen to reflect exploration of the parameter space performed in a previous paper by Destexhe (Destexhe, 1998), and were meant to be interpreted qualitatively. The default mode of the model exhibits spontaneous oscillations due to spontaneous discharges in TC neurons. After allowing the network to reach steady state, a 0.7nA stimulus was applied for 100ms to a group of five pyramidal neurons, triggering additional oscillations.

**Fig. 1.**
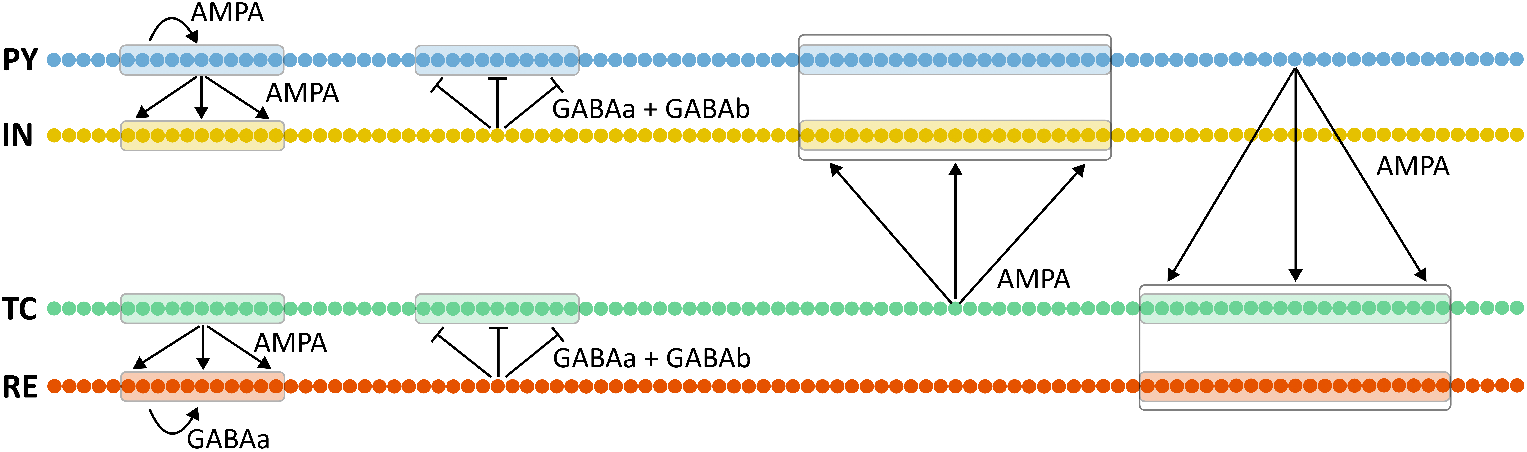
Schematic representation of the thalamocortical network. Each neuron is represented by a dot, and each layer consists of 100 cells with synaptic connectivity within and between layers as shown. Within the cortex and thalamus, each neuron connects to 11 postsynaptic neurons while between the cortex and thalamus, each neuron connects to 21 postsynaptic neurons as shown. Adapted from Destexhe et al. (Destexhe et al., 1998).

The local field potential (LFP) was calculated using the formalism by Destexhe (Destexhe, 1998), in which cortical pyramidal neurons are arranged in a single line 20*µ*m apart. The extracellular site is considered to be 50*µ*m opposite to the center of the line, and the LFP at this site was calculated from postsynaptic currents of all pyramidal neurons:

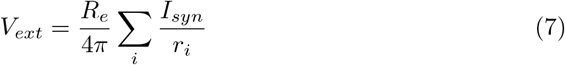

where *V*_*ext*_ is the potential at a defined extracellular site, *R*_*e*_ = 230Ω *cm* is the extra-cellular resistivity, *I*_*syn*_ is the postsynaptic current, and *r*_*i*_ is the distance between the location of the postsynaptic currents and the extracellular site.

While the original model was implemented in the NEURON simulation environment, we implemented the model in NetPyne, a Python package developed by Dura-Bernal et al. that allows for high-level specification of network connectivity of biological neuronal networks and simulations using the NEURON simulator (Dura-Bernal et al., 2019). The switch to NetPyne allows for better and easier control of accessing certain model components, make changes as necessary, and better model reproducibility, keeping in mind future directions. All of the raster plots were created using NetPyne’s *plotRaster* function. All of the spectrograms were created using the *Scipy* Python library. The LFP traces were filtered using a Butterworth filter with a cutoff frequency of 15 Hz, and power spectral density analysis was performed on the filtered LFP traces using Welch’s method.

### 2.1 Modelling the effect of genetic mutations on GABAa and T-type *Ca*^2+^ currents

Several studies have demonstrated the functional consequences of mutations of genes encoding GABAa receptor subunits, particularly GABRG2 gene. In the case of mutant receptors containing the *γ*2(R43Q) subunit, there have been studies using biotinylation of surface receptors and immunoblotting that have demonstrated reduced surface expression (Kang & Macdonald, 2004). Furthermore, electrophysiological recording studies have shown a reduced amplitude of GABAaR currents in cells containing this particular mutant receptor, consistent with decreased neuronal feedforward inhibition in the cortical circuitry (Bianchi, Song, Zhang, & Macdonald, 2002; Currie et al., 2017). The effect of the GABRG2 mutation was implemented in our model by altering the parameter denoting the maximum conductance of the GABAa current 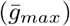 in cortical pyramidal neurons, as illustrated below:

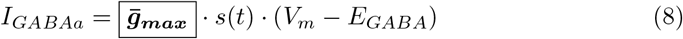

The functional consequences of a spectrum of mutations in the CACNA1H gene which encodes the Cav3.2 T-type calcium channel include a significant impact on sustained burst firing in RE neurons and the subsequent propagation of spike wave discharges (Cain et al., 2018). In particular, this includes alteration of current peak amplitude as well as activation and inactivation time (Cain et al., 2018; Vitko et al., 2005). As such, following the formalism by Knox et al. (Knox et al., 2018), we modelled the effect of CACNA1H mutations by introducing a vertical shift in the in activation time (*τ*_*h*_) of the T-type calcium current for RE neurons in our model:

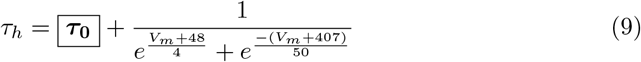

### 2.2 Modelling the effect of allopregnanolone

The effect of allopregnanolone (ALLO) was implemented on the synapse level only, focussing on GABAa-receptor mediated synapses, modelled by Equation (8). The conductance function dynamics for these synapses are described by the following equations (details of which can be found in Ermentrout and Terman (Ermentrout & Terman, 2010)):

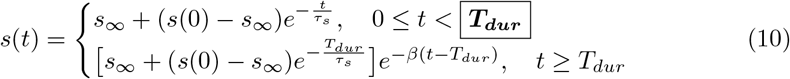

where

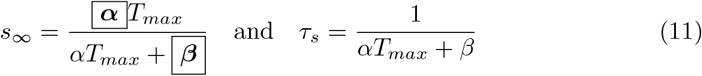

The effect of ALLO was modelled by making appropriate changes to the rise and decay time constants which describe the binding of neurotransmitter (*α* and *β* respectively) as well as the duration of neurotransmitter-mediated pulse (*T*_*dur*_), as highlighted in Equations (10)–(11). These changes were informed by fitting our synapse model to available experimental data while relying only on relative changes between control GABAa receptor activity and activity post application of ALLO (Majewska et al., 1986; Paul & Purdy, 1992). In this regard, given that the range of values for these relative changes is based on experiments conducted under different conditions, the values are meant to be interpreted qualitatively. A comparison of simulated GABAa current using Control synapses and post-ALLO synapses is shown in Figure 2 and the parameters of interest for each synapse type is given in Table 1.

**Table 1.** Rise and decay time constants, and neurotransmitter duration for Control and post-ALLO synapses given by Equations (10)–(11).

| | $\alpha$ ($(\text{ms} \cdot \text{mM})^{-1}$) | $\beta$ ($\text{ms}^{-1}$) | $T_{dur}$ (ms) |
| --- | --- | --- | --- |
| Control | 20 | 0.162 | 0.3 |
| post-ALLO | 22.87 | 0.095 | 0.75 |

**Fig. 2.**
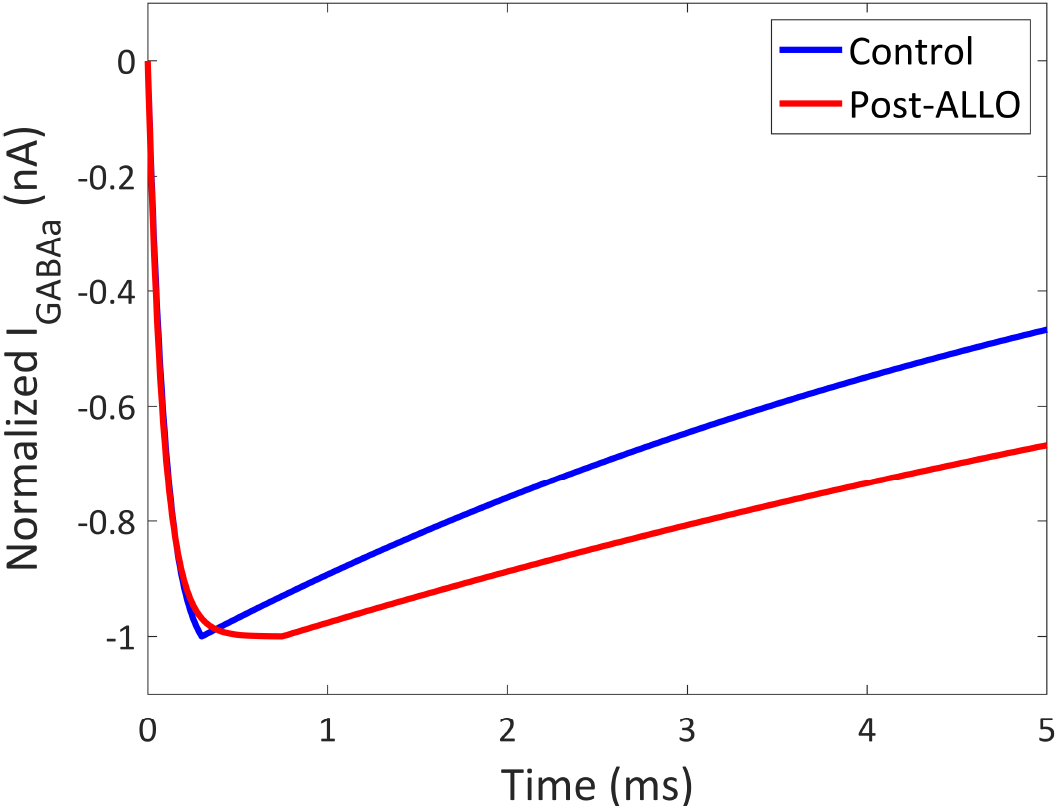
A comparison of simulated GABA current for a single spike using parameters corresponding to Control and post-ALLO synapses given by Equations (10)–(11) and Table 1.

## 3 Results

### 3.1 Baseline healthy and diseased states using control synapses

The first part of this study included modelling and characterizing baseline network behaviour that was either a healthy state exhibiting spindle oscillations (7-10 Hz) or a diseased state exhibiting spike-wave discharges (2.5-5 Hz). For our baseline simulations, all GABAa synapses were of the Control type, as described in Table 1. Network connectivity was initialized to produce a default mode of spindle oscillations with a peak frequency of 8.5 Hz, as shown in Figure 3A and 3C. The network could then be transformed to a diseased state using the two mechanisms, namely cortical disinhibition and *Ca*^2+^ conductance alteration, individually as well as together. Due to a large enough reduction of GABAaR mediated inhibition in the cortex (maximal conductance reduced to 8% of baseline), the network produces a 4.5 Hz SWD pattern as shown in Figure 3B and 3D. Similarly, an altered inactivation time of the T-type calcium current for RE neurons together with a reduction in cortical GABAa conductance resulted in the network’s ability to generate a pattern associated with SWDs, exhibiting a 3.5 Hz peak network frequency as shown in Figure 3E. In this mixed-mechanism approach, the inactivation time of the T-type calcium current was increased from 28.3 ms to 60 ms, while the cortical GABAa conductance was reduced to 25% baseline.

**Fig. 3.**
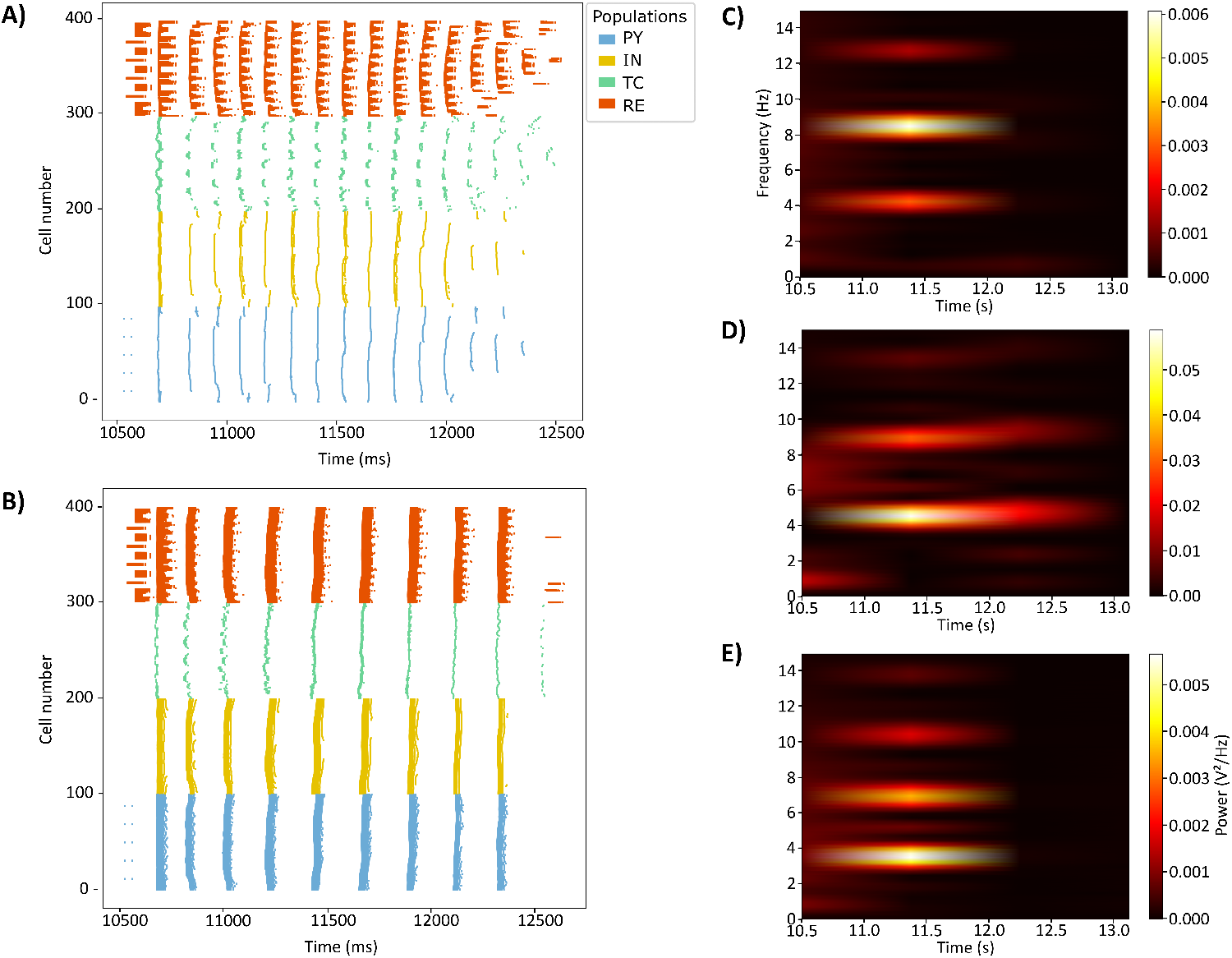
A raster plot of the full network exhibiting spindle oscillations (A) and SWD (B) behaviour. The network in (B) was simulated using the cortical disinhibition mechanism only. (C) Network behaviour corresponding to spindle oscillations exhibits maximal power at a frequency of 8.5 Hz. (D) SWD behaviour using the cortical disinhibition mechanism results in maximal power at 4.5 Hz. (E) The network displays SWD behaviour using the altered-*Ca*^2+^ conductance mechanism with maximal power at approximately 3.5 Hz. All of the simulations were performed using Control GABAa-receptor mediated synapses.

### 3.2 Allopregnanolone has an ameliorating effect on SWDs

The second part of this study involved modelling the effect of allopregnanolone by altering parameters associated with the GABAa current as described in the Methods section. The network was initialized to exhibit SWDs using both of the previously described mechanisms. The effect of ALLO was implemented on all GABAa synapses in the model, namely IN-PY, RE-TC, and RE-RE connections. It was observed that allopregnanolone has an ameliorating effect on SWDs such that the spiking frequency of the network increases and transitions from that within the range for SWDs to a range associated with spindle oscillations, as shown in Figure 4. The effect of ALLO appeared to cause a more drastic transition in network behaviour from SWDs to spindle oscillations when the underlying pathophysiology was due only to cortical disinhibition (Figure 4A and 4B). In the case of initializing the network using the altered *Ca*^2+^ conductance mechanism, the effect of ALLO was not as effective in the resolution of SWDs (Figure 4C and 4D).While the power in both frequency ranges increased with a greater increase in the range associated with spindle oscillations, the frequency range with maximum power continued to be the one for SWDs.

**Fig. 4.**
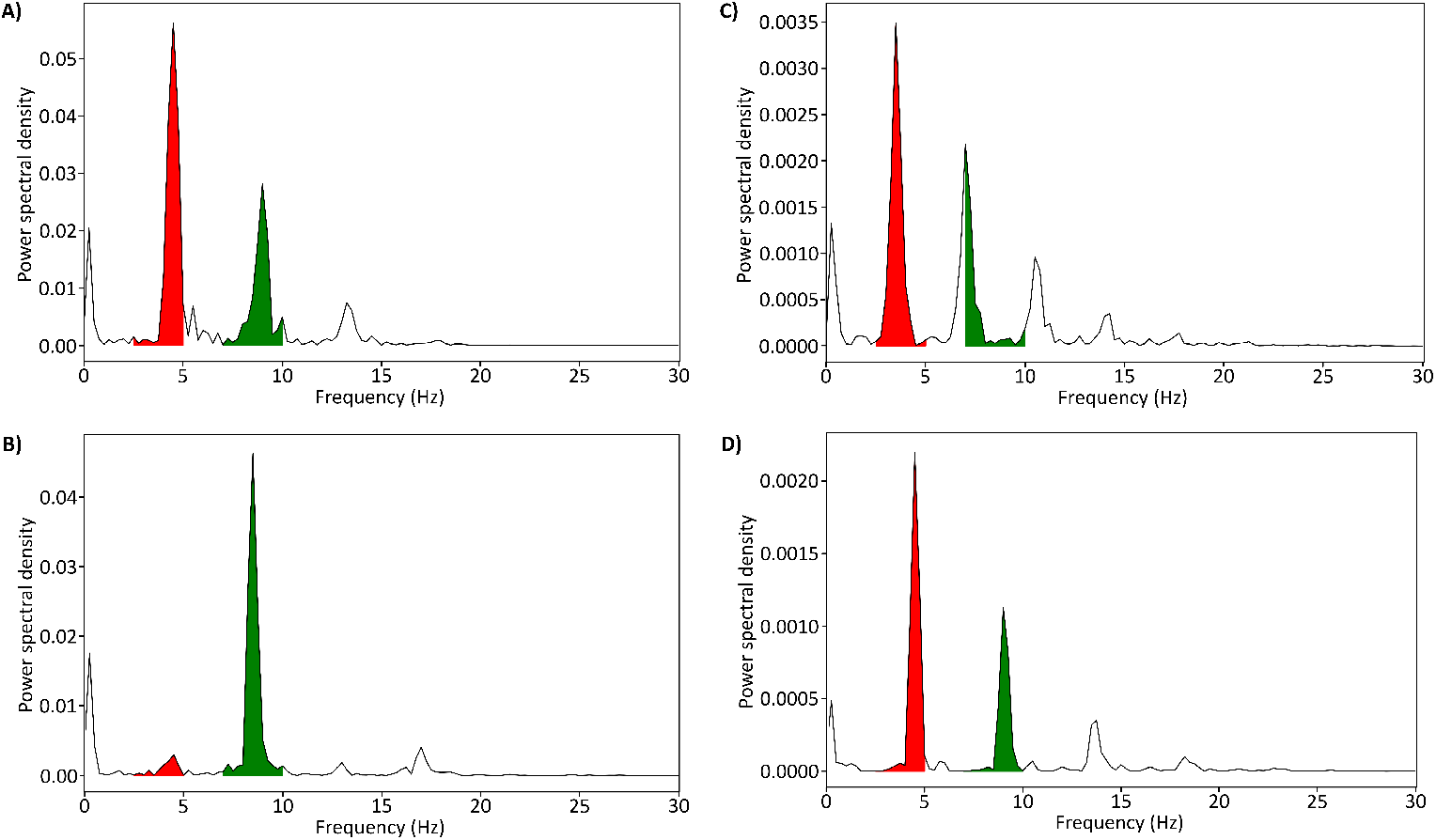
Power spectral density for network behaviour simulated using Control synapses (A and C) and post-ALLO synapses (B and D). (A)-(B): The network was initialized using the cortical disinhibition mechanism only (8% baseline cortical GABAa conductance). (C)-(D): The network was initialized using the altered *Ca*^2+^ conductance and GABAa conductance mechanism (*τ*_0_ = 60 ms and cortical GABAa at 25% of baseline). The shaded regions represent relative power in either the SWD (2.5-5 Hz; red) or Spindles (7-10 Hz; green) frequency range.

### 3.3 Exploring the selective action of allopregnanolone

It is known that neurosteroids such as allopregnanolone may be synthesized de novo in the central nervous system, and that endogenous allopregnanolone levels are brain region specific (Diviccaro, Cioffi, Falvo, Giatti, & Melcangi, 2022; Schumacher et al., 2014). We modelled these region-specific differences by exploring the selective action of ALLO on synapses between specific neuron types in the cortex and thalamus. In particular, this was implemented by making changes to parameters pertaining to the GABAa current (as described in 2.2) for the following synapses: (i) IN-PY (cortex only); (ii) IN-PY and RE-TC; (iii) IN-PY and RE-RE; (iv) RE-TC; (v) RE-RE; (vi) RE-TC and RE-RE (thalamus only).

The network behaviour in each simulation was characterized using a measure of the relative power within either the frequency range associated with SWDs or that associated with spindle oscillations (Figure 5A) as well as the peak frequency of the network (Figure 5B). Here we have defined peak network frequency as the frequency with the most power. The initial SWD state in each simulation was generated using the cortical disinhibition mechanism only. When implemented on the cortex alone, the action of ALLO makes little difference to the overall network behaviour in terms of changes in the peak frequency or the relative power in both frequency ranges. On the thalamus alone, there is a clear transition from SWDs to spindle oscillations as demonstrated by a raster plot (not shown) as well as a change in the peak network frequency from 4.5 Hz to 8.0 Hz. Exploring further within the thalamus, the action of ALLO on the RE-TC synapses alone makes little difference but paired with altered synapses in the cortex, there appears to be a very small increase in network frequency. However, the associated pattern of the network is not clearly defined spindle oscillations. Furthermore, alterations in the RE-RE synapses alone resulted in a significant change in network behaviour as demonstrated by alternating bursts in the TC population seen on the raster plot, as well as a change in peak frequency of the network from 4.5 Hz to 8.0 Hz. When paired with alterations in the cortex, this transition in peak frequency was further enhanced to 8.5 Hz.

**Fig. 5.**
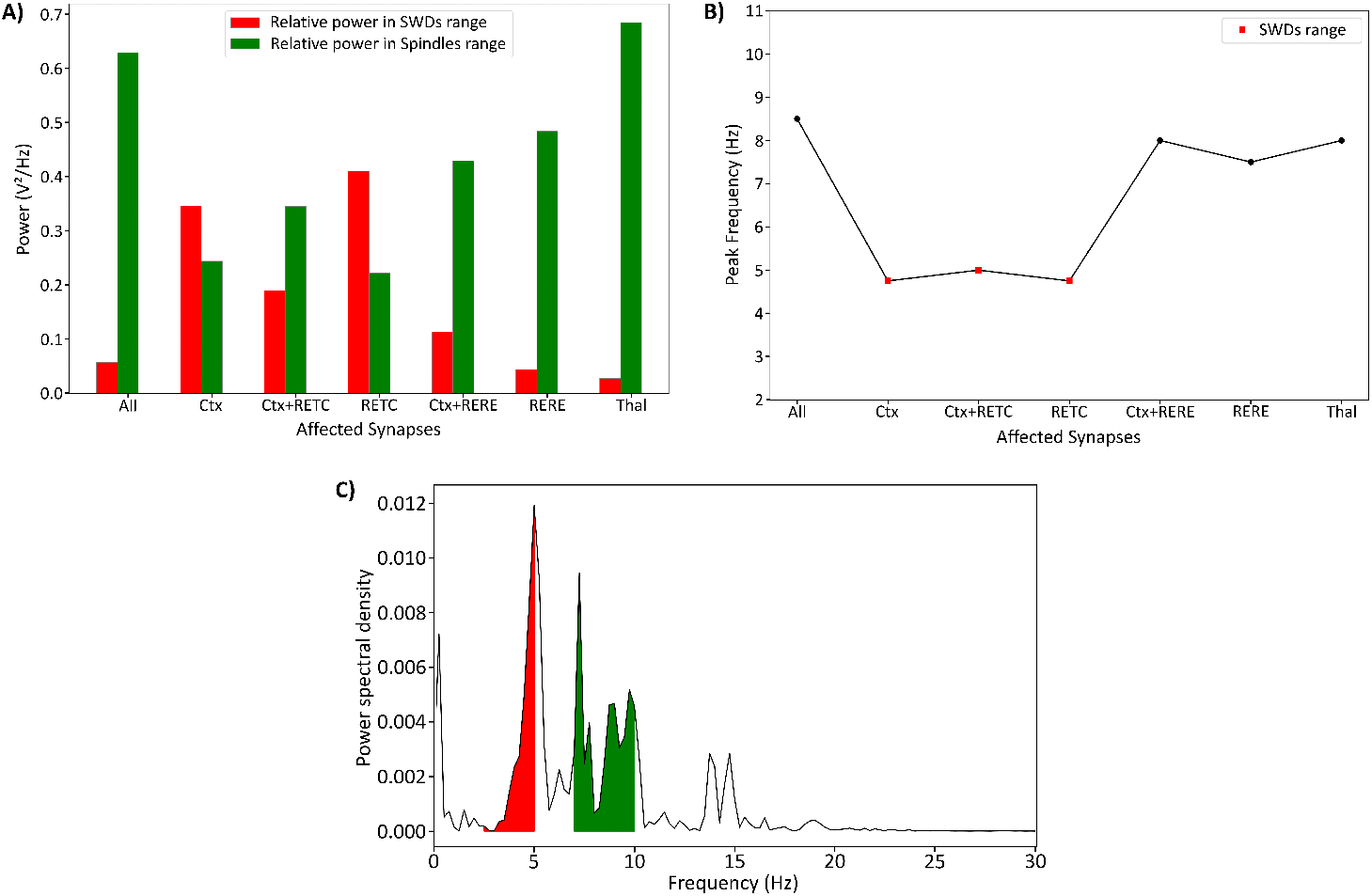
Relative power in frequency range corresponding to spindle oscillations and spike-wave discharges (A) and the peak frequency (B) for network simulations with post-ALLO synapses between specific neuron types in the cortex and thalamus. For all the simulations, the diseased state was produced using the cortical disinhibition mechanism only. (C) Power spectral density for network behaviour after implementing the effect of ALLO only on IN-PY and RE-TC synapses together.

It is to be noted that except for alterations in the IN-PY and RE-TC synapses together, all of the simulations showed consistency with regards to a relation between the relative power within a given frequency range and the peak frequency of the network. As such, more power within the range corresponding to SWDs was correlated with a peak network frequency within the same range. However, in the case of the effect of ALLO on IN-PY and RE-TC synapses together, the relative power was greater in the 7-10 Hz range corresponding to spindles, even though the peak frequency of the network was 5 Hz, within the range corresponding to SWDs (Figure 5C). This was due to a narrower peak around 5 Hz, therefore contributing towards less power as compared to the 7-10 Hz range. Overall, these results suggest that the effect of allopregnanolone is most significant in the thalamus, particularly on the RE-RE synapses which appear to be a critical contributor to the resolution of spike-wave discharges.

## 4 Discussion

In this study, we modelled the effect of allopregnanolone on a thalamocortical circuit associated with childhood absence epilepsy. Our network was initialized to a diseased, spike-wave discharge producing state that was informed by the genetic basis of the disease related particularly to GABAa-mediated inhibition and T-type calcium channel activity. The available data about the effect of hormones and neurosteroids is at two very different scales: the large-scale effect on EEG/seizures, and the small-scale effect on individual neurons/synapses. Our work is intended to fill in the gap as to how or why the in-vitro results lead to the observed EEG or network behaviour. The effect of hormones and neurosteroids on network behaviour has not been modelled previously which makes this study interesting even outside of the context of CAE.

Our results suggest that allopregnanolone has an overall ameliorating effect on absence seizures, pushing a diseased state of the network towards a healthy nonepileptic state when the disease mechanism was due to cortical disinhibition only. On the contrary, the action of ALLO appeared to be insufficient to counteract the aggravating effect of altered T-type *Ca*^2+^ channel activity contributing to sustained SWDs. As a result, our investigation of ALLO and its interaction within the thalamocortical circuit was limited to the cortical disinhibition mechanism only. Exploring the role of ALLO further on individual synapses comprising the GABAergic system revealed the selective action of this neurosteroid in either sustaining SWDs or resolving them. The importance of the GABAergic system in the pathophysiology of epileptic seizures has existed for a long time and is no different in the case of CAE. Our results are also consistent with the brain region-specific function of GABA across the thalamocortical network in the context of absence seizures. In the cortex, a hypofunction of GABAergic neurons is shown to aggravate SWDs (D’antuono et al., 2006; Luhmann, Mittmann, van Luijtelaar, & Heinemann, 1995). On the contrary, in the thalamus, a hyperfunction of GABA transmission in the thalamocortical neurons is observed to increase SWD duration while increased GABA in the thalamic reticular neurons suppresses SWDs (Liu, Vergnes, Depaulis, & Marescaux, 1991; Pinault et al., 1998). The prolonged action of GABA as a consequence of the effect of ALLO in TC neurons resulting in a continued SWD pattern can be correlated with increased SWD duration, or at least as a lack of positive impact on SWD behaviour. Similarly, implementing ALLO-affected synapses in RE neurons very clearly resulted in the suppression of SWD activity in our network. It is interesting to note this consistency in the results considering that the mechanism by which there is increased GABA-mediated activity is different. As such, our approach involves implementing the effect on the intricacies of the binding process of GABAa receptor function via time constants and neurotransmitter duration which is different from the classical approach of a change in channel conductance which is associated with channel blocking or up/down regulation. This provides a new avenue to explore function of GABAa receptors with regards to computational modelling in other contexts.

One pertinent question that arises is, if the application of progesterone or its metabolite allopregnanolone, as suggested by clinical evidence in humans and animals, aggravates spike-wave discharges, then are our results contradictory in that they suggest an ameliorating effect by allopregnanolone? We do not believe this to be the case as it is possible that the available experimental data does not provide the full picture related to remitting absence seizures. It is important to note that most of the clinical evidence involves studying ‘non-resolving’ absence epilepsy models such as the WAG/Rij or GAERS rodent models. That is, these animal models exhibit absence seizures throughout their lifetime, and do not exhibit remission at any period whatsoever. On the contrary, the intrinsic connectivity and simplicity of our model corresponds to one describing ‘resolving CAE’. Therefore, it is possible that ALLO’s remedying effect on SWDs is only true for resolving CAE networks. In fact, there are studies that have shown there exist pre-treatment intrinsic connectivity differences between patients who ultimately experience remission and those who do not (Tenney et al., 2018). There is also evidence of enhanced extrasynaptic GABAa-receptor mediated (tonic) inhibition in thalamocortical neurons in the GAERS model (Cope et al., 2009). Interestingly, allopregnanolone is one of the most potent extrasynaptic receptor modulators (Meltzer-Brody & Kanes, 2020). Both of these differences are not captured in our existing model. This leads us to hypothesize that remission occurring during adolescence in “most cases” could be associated with the action of allopregnanolone bringing a beneficial change only to individuals that are predisposed to remission as a result of either intrinsic connectivity differences or differences in tonic inhibition. Our future directions include exploring some of this very aspect, as it was one of the major limitations of this present work. In particular, this would involve including greater detail in the cortical component of the model to include different brain regions, especially the frontal and parietal cortex. Furthermore, it would be useful to explore other mechanisms by which a diseased state can be achieved, especially mutations in genes encoding other ion channels such as sodium and potassium channels, as well as those encoding specific GABAa receptor subunits, and how these different mechanisms interact with the effect of ALLO.

## 5 Conclusion

Understanding the role of sex hormones in the resolution of CAE may explain why remission does not occur in nearly a quarter of cases. Results from such research can therefore lead to better informed therapeutic decisions relating to early interventions such as hormonal therapy and/or antiepileptic drug treatment, which can be tailored to intrinsic network connectivity arrangements, perhaps based on functional connectivity analysis of neuroimaging data of patients.

## Author Contributions

Each author has substantially contributed to conducting this study and in drafting this manuscript.

## Acknowledgments

We would like to thank Dr. Andrew Knox and Dr. Eugenio Urdapilleta for their assistance/clarification with the model and use of NetPyne.

## Model availability

Upon publication, the model code will be made publicly available on ModelDB.

